# A Novel In Vitro Potency Assay Demonstrating the Anti-Fibrotic Mechanism of Action of CDCs in Deramiocel

**DOI:** 10.1101/2025.09.18.675440

**Authors:** Yujia Li, Justin B. Nice, Marya Kozinova, Stephanie Adachi, Linda Marbán, Kristi Elliott, Minghao Sun

## Abstract

Duchenne muscular dystrophy (DMD) is characterized by progressive skeletal and cardiac muscle degeneration driven by inflammation and fibrosis, ultimately leading to cardiomyopathy and premature death. Deramiocel, an allogeneic cell therapy composed of cardiosphere-derived cells (CDCs), has demonstrated potent anti-fibrotic and immunomodulatory effects in preclinical models and clinical trials, including HOPE-2 and its open-label extension (HOPE-2 OLE), where Deramiocel treatment significantly attenuated progression of skeletal and cardiac muscle dysfunction. Here, we describe the development of a novel in vitro potency assay to quantify the anti-fibrotic activity of Deramiocel. Conditioned media (CM) from multiple Deramiocel manufacturing lots significantly suppressed expression of collagen type I alpha 1 (COL1A1) and collagen type III alpha 1 (COL3A1) in primary human dermal fibroblasts compared with non-conditioned media controls, establishing a robust, reproducible readout of anti-fibrotic activity. The effect was dose-dependent and abrogated by sequential depletion of exosomes and soluble proteins, implicating both as critical mediators of Deramiocel’s mechanism of action. Importantly, CDCs in deramiocel lots classified as potent by this assay were shown to exert a clinically meaningful benefit in DMD patients in the HOPE-2 and HOPE-2 OLE studies. This assay represents a mechanistically informative, therapeutically relevant, reproducible, scalable, and regulatory-compliant approach for assessing Deramiocel potency, enabling consistent manufacturing and facilitating the continued development of Deramiocel as a disease-modifying therapy for DMD.

## 1. Introduction

Duchenne muscular dystrophy (DMD) is a severe, X-linked disease caused by mutations in the dystrophin gene resulting in progressive skeletal and cardiac muscle degeneration. The pathophysiology of DMD is such that the absence of dystrophin leads to accelerated injury to muscle cells resulting in a continuous cycle of muscle breakdown followed by inflammation and fibrosis. Early fibrosis is subsequently replaced by fibrofatty scar tissue as the disease progresses leading to both skeletal and cardiac muscle dysfunction as shown in Figure 1 [1]. This fibrofatty, non-contractile scarring of the myocardium leads to progressive weakening of the heart muscle, in particular the left ventricle, resulting in ever worsening cardiomyopathy. Cardiomyopathy is a universal feature of DMD and is the leading cause of death in DMD [2]. Thus, while new therapies have focused on restoration of dystrophin to address the primary disease, it is equally important to treat the inflammation and fibrosis that occurs in conjunction with the constant breakdown of muscle to preserve muscle function.

**Figure 1:**
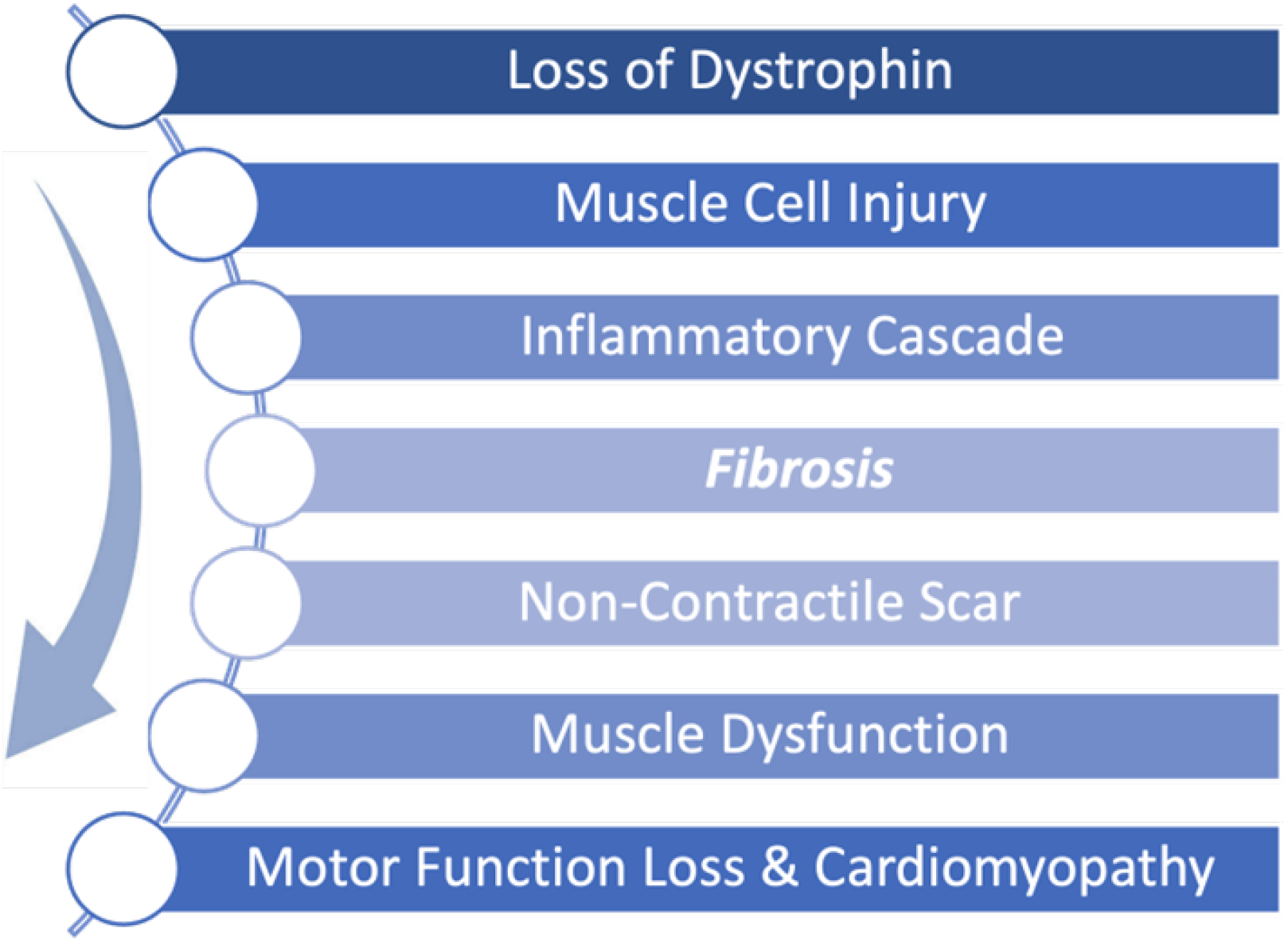
Pathophysiological progression of Duchenne muscular dystrophy (DMD) begins with a genetically caused deficiency in dystrophin expression. During normal muscle use, the absence of dystrophin renders muscle cells unable to recover from mechanical stress, leading to repeated cellular injury. This triggers an inflammatory cascade that promotes fibrosis and scar formation. Over time, non-contractile scar tissue replaces functional muscle, driving progressive muscle dysfunction, motor function loss, and cardiomyopathy.

Deramiocel is a human allogeneic cell therapy containing cardiosphere-derived cells (CDCs) derived from heart tissue which has been shown in numerous publications to have anti-fibrotic and immunomodulatory activity, which in turn aids in muscle survival and regeneration [3, 4, 5, 6]. CDCs were originally developed as a therapeutic agent for myocardial infarction and cardiac dysfunction [7]. In early clinical trials focused on myocardial infarction, the efficacy of CDCs was demonstrated by a reduction in fibrotic scar size in infarcted hearts using contrast-enhanced cardiac MRI [8].

Evidence of the efficacy and bioactivity of CDCs to treat DMD first arose in several nonclinical studies (unpublished; [9]) using the *mdx* mouse model which mimics the skeletal, respiratory, and cardiac dysfunction in DMD [10, 11]. In these studies, CDCs were shown to improve skeletal muscle function as demonstrated by improved exercise capacity. Importantly, CDC treatment improved global cardiac function, similar to efficacy demonstrated in preclinical models and early clinical trials. Analysis of ejection fraction (EF) in the *mdx* mouse model after treatment with CDCs showed that EF improved within 3 weeks of treatment, driven largely by improvements in end-systolic volume (unpublished; [9]). In contrast, vehicle-treated mice continued to decline. The overall results of the studies showed that the baseline EF was sustained in CDC-treated mice (2 administrations) over a 6-month period. Importantly, a statistically significant reduction in myocardial fibrosis was observed in CDC-treated *mdx* mice as evidenced by reductions in both histological fibrosis and collagen content in the hearts of mice treated with CDCs. This reduction in fibrosis observed *in vivo* in CDC-treated hearts clearly demonstrates the anti-fibrotic mechanism of action of CDCs, aligning with published literature.

Data from the first clinical trial in DMD, the HOPE-Duchenne clinical trial, provided further evidence that CDCs, the active ingredient in Deramiocel, have anti-fibrotic activity as shown by a reduction in cardiac scarring and overall improvement in both cardiac and skeletal muscle function in DMD patients. Likewise, data from the HOPE-2, phase 2 randomized double-blind and placebo-controlled clinical trial, published in *The Lancet*, demonstrated that DMD patients treated with Deramiocel over 12 months experienced significant attenuation of disease progression. Skeletal muscle function, as measured by mid-level performance of upper limb (PUL), showed a 71% slowing of decline (*p* = 0.014). Importantly, the HOPE-2 trial showed significant preservation of cardiac function across multiple cardiac endpoints, including a 107% slowing of cardiac disease progression as measured by left ventricular ejection fraction (LVEF) (*p* = 0.002). The preservation of cardiac function with Deramiocel treatment is likely a result of the reduction in inflammation and fibrosis imparted by CDCs which in turn results in reduced non-contractile scarring of the myocardium. Together, these findings highlight the clinically meaningful and statistically significant therapeutic benefit of Deramiocel [2, 12].

Following the HOPE-2 trial, 13 patients continued on open-label treatment with Deramiocel for up to 48 months. The therapeutic benefit of Deramiocel was maintained in both cardiac and skeletal muscle function. Skeletal muscle outcomes, assessed by PUL 2.0, showed a consistent reduction in disease progression: the mean decline was 1.8 points in Year 1, 1.2 points in Year 2, 1.1 points in Year 3, and 0.6 points in Year 4. Among 10 patients with centrally read cardiac MRI assessments, the median change in LVEF after 48 months was strikingly only −0.5%, clearly demonstrating the long-term cardiac stability of DMD patients treated with Deramiocel. These findings provide strong evidence that long-term Deramiocel therapy can slow the progression of both skeletal and cardiac muscle disease in patients with DMD. The stabilization of cardiac function in DMD patients for over 4 years is unprecedented and provides an opportunity for improved quality of life and decreased mortality in DMD patients.

Exosomes isolated from CDCs have been shown to be necessary and sufficient for the bioactivity of CDCs. In fact, CDC exosomes alone were able to reproduce improvements in cardiac and skeletal myopathy observed with CDCs using *mdx* mice [13]. CDC exosomes have been demonstrated to track to sites of injury and to be rapidly taken up by cardiac and skeletal muscle cells following injury in multiple mouse studies (unpublished, [13]). CDC exosome treatment improved contractile force in *mdx* soleus muscles, reduced interstitial fibrosis, and preserved muscle architecture. Importantly, treatment with CDC exosomes reduced myocardial collagen content, reduced myocardial scarring and improved LVEF to the same levels as CDC treatment alone [14, 15, 16]. Consistent with these effects, collagen I and III levels were reduced in CDC exosome–treated *mdx* hearts [17]. In addition, the improvements in LVEF in the mouse model observed after CDC treatment were attenuated when exosome secretion was inhibited [14]. Similar therapeutic benefits have been observed in large-animal models, including reduced scarring, anti-fibrotic effects, and improved LVEF [16]. Mechanistically, this finding suggests that exosomes secreted by CDCs play a critical anti-fibrotic role in stabilizing muscle function through suppression of collagen expression [18].

Similarly, paracrine signaling via growth factor secretion by CDCs has been implicated in the immunomodulatory and anti-fibrotic mechanism of action of CDCs in published literature. CDCs have been shown to secrete large amounts of VEGF, HGF, TIMP-2, IGF-1, SDF-1, angiopoietin 2 and bFGF [19, 20]. In particular, VEGF reduces ischemia-driven fibroblast activation and plays an indirect yet significant anti-fibrotic role; HGF inhibits TGF-*β*–induced fibroblast-to-myofibroblast transition and collagen deposition and acts as a potent anti-fibrotic cytokine; and TIMP-2, which, in combination with matrix metalloproteinases (MMPs), helps maintain a balanced extracellular matrix, supports scar remodeling, and prevents excessive collagen cross-linking [21]. Additionally, other factors such as IL-10 and IGF-1 were identified as indirect suppressors of fibrosis through their anti-inflammatory and pro-survival effects, respectively. The signaling of all of these secreted factors aligns with the anti-fibrotic activity of Deramiocel, suggesting that the soluble factors secreted by Deramiocel also play a critical role in its mechanism of action.

In this paper, an *in vitro* cell-based assay was developed and subsequently validated using conditioned media (CM) from Deramiocel to assess its anti-fibrotic activity. Our findings demonstrate that Deramiocel exerts quantifiable anti-fibrotic activity *in vitro*, mediated through exosome and soluble factors in a dose-dependent manner. The anti-fibrotic activity observed *in vitro* using this assay aligns with data from non-clinical pharmacodynamic studies and importantly the clinical trial results observed in HOPE-2 and HOPE-2 OLE. The validated cell-based anti-fibrosis assay employed herein offers a controlled and reproducible platform enabling a mechanistically informative assessment of Deramiocel’s potency.

## 2. Materials and Methods

### 2.1. Cell Culture

Primary human dermal fibroblasts (HDFs) and human cardiac fibroblasts (HCFs; two donors) were obtained from PromoCell and ATCC, respectively. ATCC media to grow HDF and HCF were prepared according to manufacturer’s recommendation (ATCC/PCS-201-030 and ATCC/PCS-201-041). Cells were maintained at 37 ^*°*^C in a humidified atmosphere containing 5% CO_2_. For Deramiocel cells, they are cultured in our proprietary culture media with 20% FBS and were maintained at 37 ^*°*^C with 5% CO_2_ and 5% O_2_ with humidity.

### 2.2. Preparation of Conditioned Media

Human CDCs were thawed and seeded at 0.5 *×* 10^6^ cells per T75 flask in standard growth medium. After 24 h, the medium was aspirated, cells were washed once with DPBS (Gibco/14040-133) and 22 mL of fresh culture medium was added. Flasks were incubated for 72 h, after which supernatant was collected and clarified by centrifugation at 300 *×* g for 5 min followed by 3,000 *×* g for 15 min. The resulting supernatant was filtered through a 0.22 *µ*m Steriflip filter (ThermoFisher Scientific/569-0020), aliquoted into 15 mL tubes, and stored at −80 °C until use.

### 2.3. Preparation of Non-Conditioned Media with Varying FBS Concentrations

To generate non-conditioned control media, CDC growth medium was prepared with 5%, 10%, 15%, or 20% FBS while keeping all other components identical. Media were clarified by centrifugation at 300 *×* g for 5 min and 3,000 *×* g for 15 min, filtered through a 0.22 *µ*m filter (ThermoFisher Scientific/569-0020), aliquoted, and stored at −80 °C. Non-conditioned media with 20% FBS serves as a control in each assay.

### 2.4. Positive Control for Anti-Fibrosis Assay

Recombinant human EGF (rh EGF) was used as a positive control to confirm assay functionality. rh EGF from the Endothelial Cell Growth Kit-BBE (ATCC/PCS-100-040) was diluted 1:500 into non-conditioned medium immediately prior to use.

### 2.5. Treatment of Human Dermal Fibroblasts (HDFs)

HDFs were seeded at 2.5 *×* 10^4^ cells per well in 24-well plates and cultured overnight. The following treatment groups were applied: (i) Deramiocel CM, (ii) non-conditioned control medium, and (iii) CM from human cardiac fibroblasts. For dose-response experiments, Deramiocel CM was serially diluted (1:2, 1:5, 1:100) in non-conditioned medium. rh EGF was included in selected experiments as a positive control.

### 2.6. Exosome Depletion and Filtration Studies

To dissect the contribution of exosomes and soluble protein factors to the anti-fibrotic activity of Deramiocel CM, sequential filtration was performed. CM from two independent manufacturing lots (LOT-0193 and LOT-0196) was processed using Amicon Ultra centrifugal filters (MilliporeSigma/UFC8010 and UFC8100) with molecular weight cut-offs of 100 kDa and 10 kDa. For exosome depletion, CM was passed through a 100 kDa filter, and both the retentate (enriched in exosomes) and flow-through (exosome-depleted fraction) were collected and tested for biological activity in anti-fibrosis assays. For further resolution, CM was sequentially filtered through 100 kDa and 10 kDa cut-offs, yielding a 100 kDa retentate (exosome-enriched), a 100 kDa flow-through (exosome-depleted), and a 10 kDa flow-through (depleted of both exosomes and most soluble proteins). Each fraction, along with unprocessed CM and non-conditioned medium, was evaluated in the anti-fibrosis assay, with rh EGF included as positive control.

### 2.7. RNA Extraction and qRT-PCR

After 72 h of treatment, total HDF RNA was extracted using the RNeasy Micro Kit (Qiagen/74004 and 79656) according to the manufacturer’s instructions. RNA concentration and purity were measured using a NanoDrop spectrophotometer (Thermo Fisher). To proceed with cDNA synthesis the RNA concentration must be ≥20 ng/*µ*L and the A260/A280 between 1.8–2. Complementary DNA (cDNA) was synthesized from RNA using the iScript cDNA Synthesis Kit (Bio-Rad/1708891) according to the manufacturer’s instructions. qRT-PCR was performed using TaqMan Gene Expression Mastermix (Applied Biosystems/4369016) on a QuantStudio 7 Flex Real-Time PCR System (Thermo Fisher). qPCR probes for COL1A1 (FAM), COL3A1 (FAM) and GAPDH (VIC) were ordered (Thermo Fisher). GAPDH was used as a housekeeping control. Relative expression levels were calculated using the 2-ΔΔCt method against NCM. As part of the assay qualification and validation, qPCR suitability criteria were established. A standard curve must show a slope of -3.2 to -3.9, technical triplicates must have a SD ≤0.5 (one may be removed from analysis). No-template control samples must have a Cq ≥ 35/undetermined. Biological triplicates must have a standard deviation of Cq≤20%. Lastly, the positive control (rh EGF) must show ≤50% relative gene expression for COL1A and COL3A.

### 2.8. LEGENDplex Flow Assay

Further characterization of secreted factors was done using the LEGENDplex^™^ Human Angiogenesis Panel 1 (Biolegend/741214). Non-conditioned media and conditioned media from three Deramiocel lots were incubated per the manufacturer’s instructions undiluted and diluted 1:1. The beads were gated based on scattering and the fcs files were analyzed using the LEGENDplex^™^ online software. The level of expression of the secreted factors in Deramiocel conditioned media was normalized to non-conditioned media.

### 2.9. Statistical Analysis

All assays were performed in biological triplicates, with data expressed as mean *±* standard deviation (SD). Statistical analyses were conducted using GraphPad Prism 9.1. Comparisons between groups were performed using one-way ANOVA followed by post-hoc correction for multiple comparisons or 2-tailed t-tests as needed. p-values *<* 0.05 were considered statistically significant, and significance levels were denoted as follows: **** p *<* 0.0001; *** p *<* 0.0005; ** p *<* 0.001; * p *<* 0.05; ns=not significant.

For the established assay, the historical performance of up to 32 HOPE-2 and HOPE-2 OLE clinical lots (with up to 36 months of clinical data at the time of analysis) and up to 58 additional lots tested across as many as five fibroblast working cell banks derived from a single donor was analyzed to set the acceptance criteria for the antifibrotic potency assay. COL1A and COL3A inhibition values were log_2_-transformed and evaluated for normality. Data from the working cell banks were combined using weighted means and standard deviations to generate donor-specific thresholds, with acceptance criteria defined as the pooled mean minus three standard deviations for each marker (COL1A and COL3A). For presentation purposes, these inhibition-based thresholds were converted to equivalent relative expression levels.

## 3. Results

### 3.1. Anti-fibrotic Activity of CDCs in Deramiocel Correlates with Preclinical and Clinical Potency

To evaluate the anti-fibrotic activity of CDCs, an in vitro assay was developed using a co-culture system of fibroblasts with conditioned media (CM) collected from CDCs in Deramiocel (Figure 2). Briefly, CDCs in Deramiocel were cultured to generate CM enriched with secreted exosomes and factors, which was subsequently applied to primary human dermal fibroblasts (HDFs). HDFs were selected as the reporter cell line because fibroblasts are the principal effector cells in fibrosis and dermal fibroblasts provide a reproducible, well-characterized, and scalable system for in vitro testing. Following co-culture, expression of collagen type I alpha 1 (COL1A1) and collagen type III alpha 1 (COL3A1) was measured by qRT-PCR. These genes were chosen as readouts because they represent the major extracellular matrix proteins deposited during fibrotic remodeling and are widely recognized biomarkers of fibroblast activation. Non-conditioned media serves as a control in the assay. Reduction in COL1A and COL3A expression therefore provides a direct and clinically relevant measure of the anti-fibrotic activity of Deramiocel.

**Figure 2:**
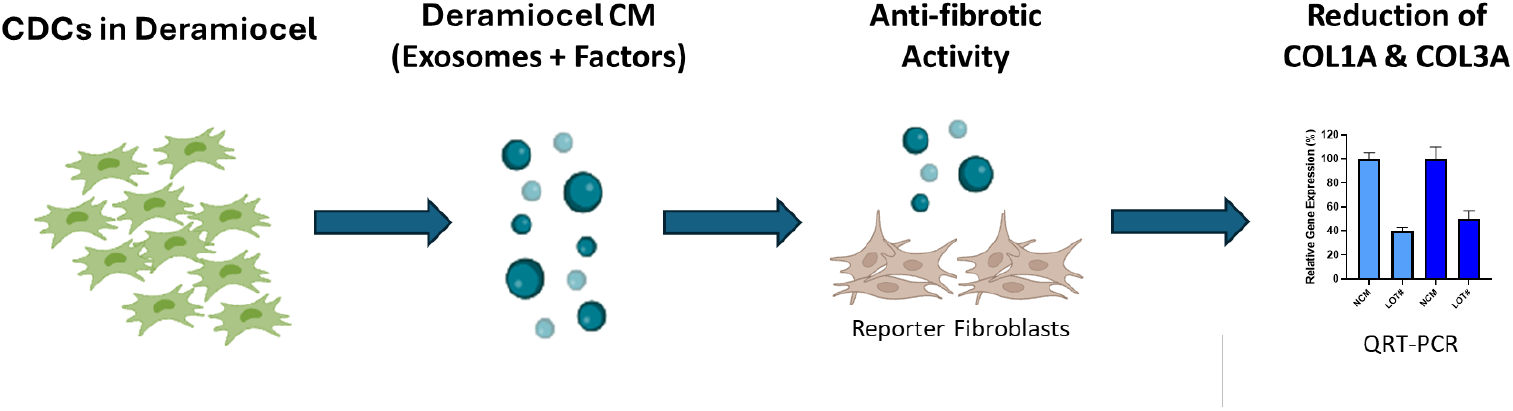
Workflow of the anti-fibrosis potency assay. An in vitro assay was developed using a co-culture system of fibroblasts with conditioned media (CM) collected from CDCs in Deramiocel. Following co-culture, expression of collagen type I alpha 1 (COL1A1) and collagen type III alpha 1 (COL3A1) was measured by qRT-PCR.

The bioactivity of three Deramiocel lots, LOT-0121 (utilized in the HOPE-2 trial), LOT-0144 (utilized in the HOPE-2 open-label extension [OLE]), and LOT-0100 (not yet tested clinically), was initially assessed in the anti-fibrosis assay. As shown in Figure 3, conditioned media collected from all three lots of Deramiocel significantly reduced the expression of both COL1A and COL3A compared with the non-conditioned media control (NCM). Notably, the two lots employed in HOPE-2 and HOPE-2 OLE were classified as potent both in the anti-fibrosis assay and in the mouse myocardial infarction (MI) model, and were also demonstrated to be clinically effective. Thus, the anti-fibrotic activity as demonstrated by the reduction in collagen expression observed with these two deramiocel lots is consistent with potency classified by the in vivo MI mouse model and with clinical potency.

**Figure 3:**
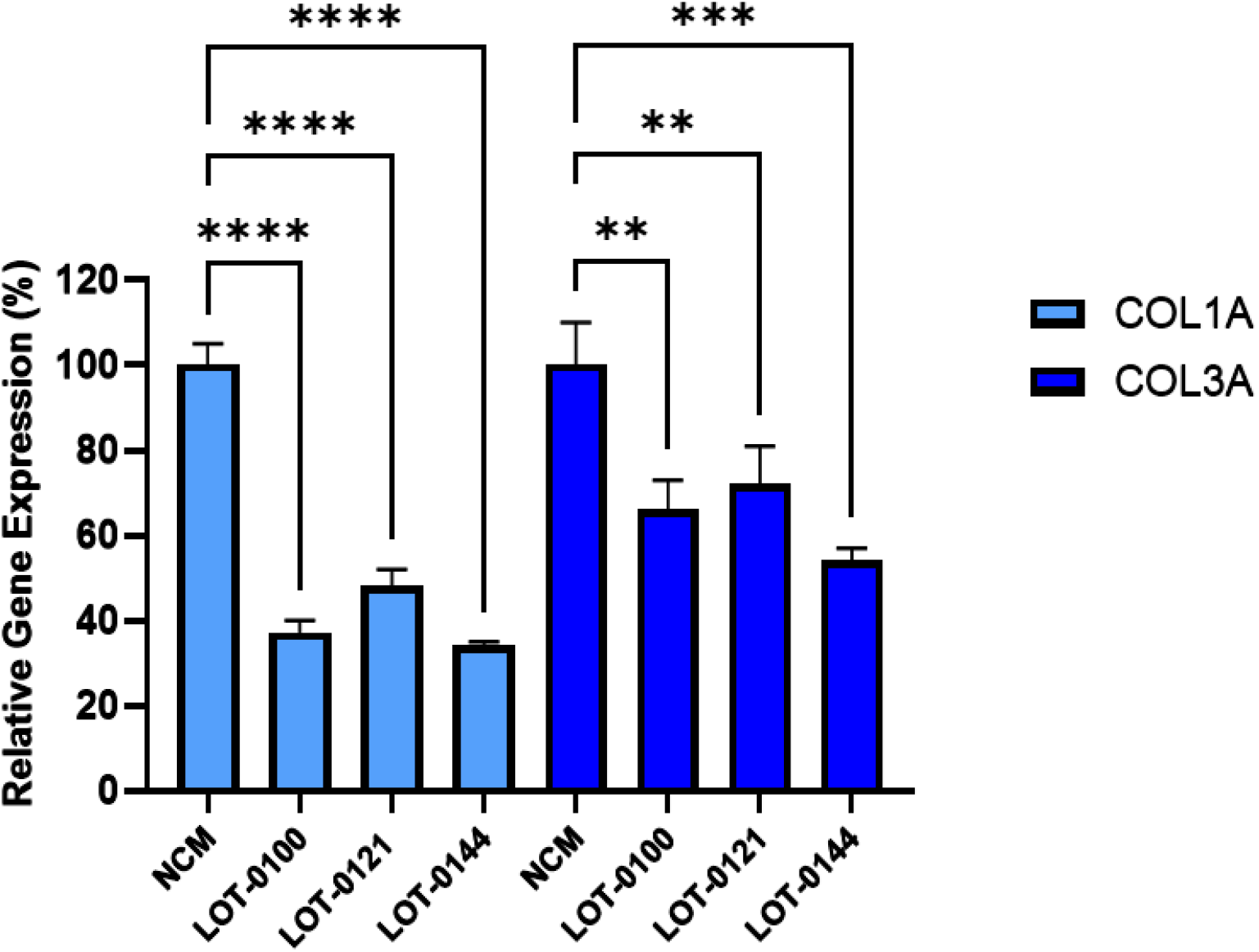
Initial testing of three independent Deramiocel lots. Significant reduction of collagen gene expression (COL1A, COL3A) was observed in fibroblasts following co-culture with Deramiocel conditioned media (CM) compared to non-conditioned media (NCM) control (set at 0%, no inhibition). Assays were performed in triplicate, and data are presented as mean ± SD, normalized to NCM. Statistical significance: **** p<0.0001, *** p<0.0005, ** p<0.001.

Building on these findings, 11 additional lots of Deramiocel were subsequently evaluated, including LOT-0144 (used in HOPE-2 OLE; repeated for reproducibility), LOT-0124 (used in HOPE-2), LOT-0139 (used in HOPE-2 OLE), and LOT-0147 (used in HOPE-2 OLE). As shown in Figure 4, CM from all 11 lots significantly decreased COL1A (left panel) and COL3A (right panel) expression relative to the non-conditioned media (NCM) control. Importantly, results obtained with LOT-0144 were reproducible in this independent study, further demonstrating the reproducibility of the assay. Consistent with earlier observations, the four lots used clinically in HOPE-2 or HOPE-2 OLE were classified as potent in the mouse MI model and were shown to effective clinically. Collectively, the anti-fibrotic activity as demonstrated by the reduction in collagen expression induced by these four deramiocel lots is consistent with potency classified by the in vivo mouse MI model and with clinical potency.

**Figure 4:**
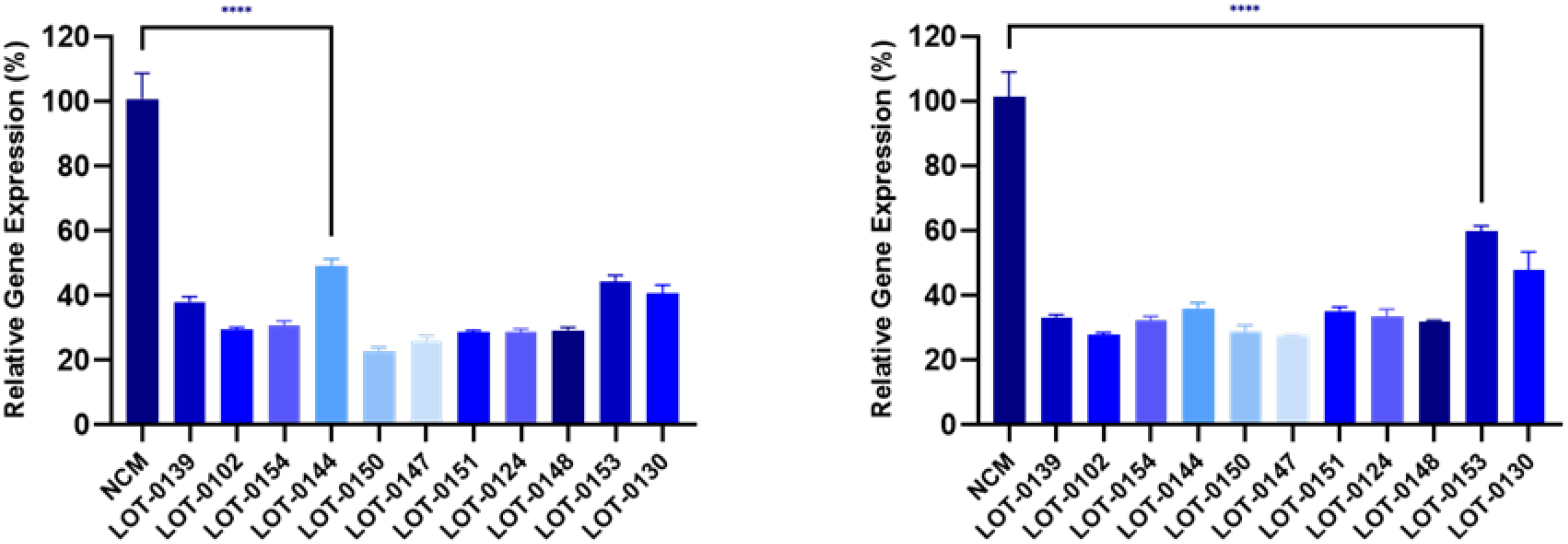
Reduction of collagen gene expression (COL1A, left panel; COL3A, right panel) in fibroblasts after co-culture with CM from 11 Deramiocel lots compared to NCM control (Set as 0%, no inhibition). Assays were performed in triplicate, and data are presented as mean ± SD, normalized to NCM. A representative statistical comparison is shown for the lot with the least reduction in collagen expression relative to NCM (**** p<0.0001).

To confirm data presented in Figure 4, independent vials of deramiocel were thawed and CM collected for evaluation in an independent anti-fibrosis assay (data not shown). CM collected from all repeat lots induced a statistically significant reduction in both COL1A and COL3A expression when compared to the non-conditioned media (NCM) control; demonstrating that the assay and data are consistent and reproducible.

### 3.2. Anti-fibrotic Effect is Attributed to Deramiocel Secreted Factors

To ensure that the anti-fibrotic activity observed in these studies was not due to protein or nutrient depletion in the conditioned media, the effect of reduced fetal bovine serum (FBS) concentrations was evaluated to mimic potential nutrient loss in CDC media. As shown in Figure 5, decreasing FBS concentrations from 20% to 5% in non-conditioned CDC media (NCM) resulted in increased expression of both COL1A and COL3A, as measured by qRT-PCR. These findings indicate that the reduction in collagen expression is not caused by protein or nutrient loss. Furthermore, the data supports the conclusion that the anti-fibrotic effect is primarily mediated by bioactive factors secreted by CDCs in Deramiocel, such as exosomes and growth factors.

**Figure 5:**
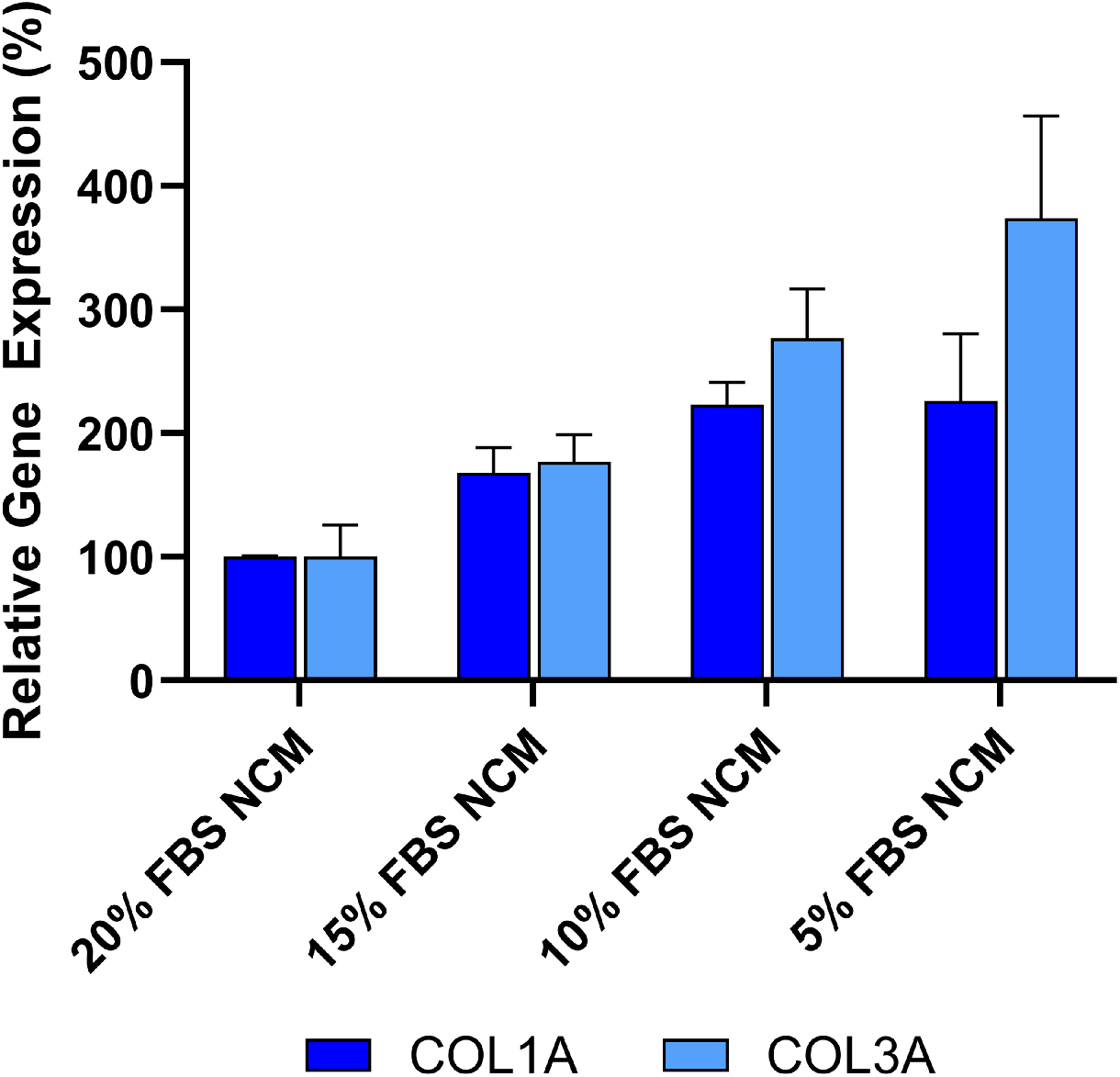
Anti-fibrotic effect is not caused by protein or nutrient depletion from non-conditioned media (NCM). Fibroblasts were cultured with NCM containing decreasing concentrations of FBS to mimic nutrient loss. Assays were performed in triplicate, and data are presented as mean ± SD, normalized to 20% NCM.

### 3.3. Assay Specificity Demonstrated Using Non-Deramiocel Cells

To evaluate the specificity of the assay, conditioned media (CM) from non-CDC cells was tested. Human cardiac fibroblasts (HCF) were selected to evaluate in the assay since they are also derived from human hearts. Note that other cells from the heart could not be evaluated in the assay due to the inability of those cells to grow in CDC media required for the assay. As shown in Figure 6, CM from primary HCFs derived from two donors did not inhibit COL1A or COL3A expression relative to the non-conditioned media (NCM) control, as assessed by qRT-PCR. Accordingly, these HCFs would be classified as non-potent due to the absence of anti-fibrotic activity, supporting the assay’s specificity for CDC-mediated effects.

**Figure 6:**
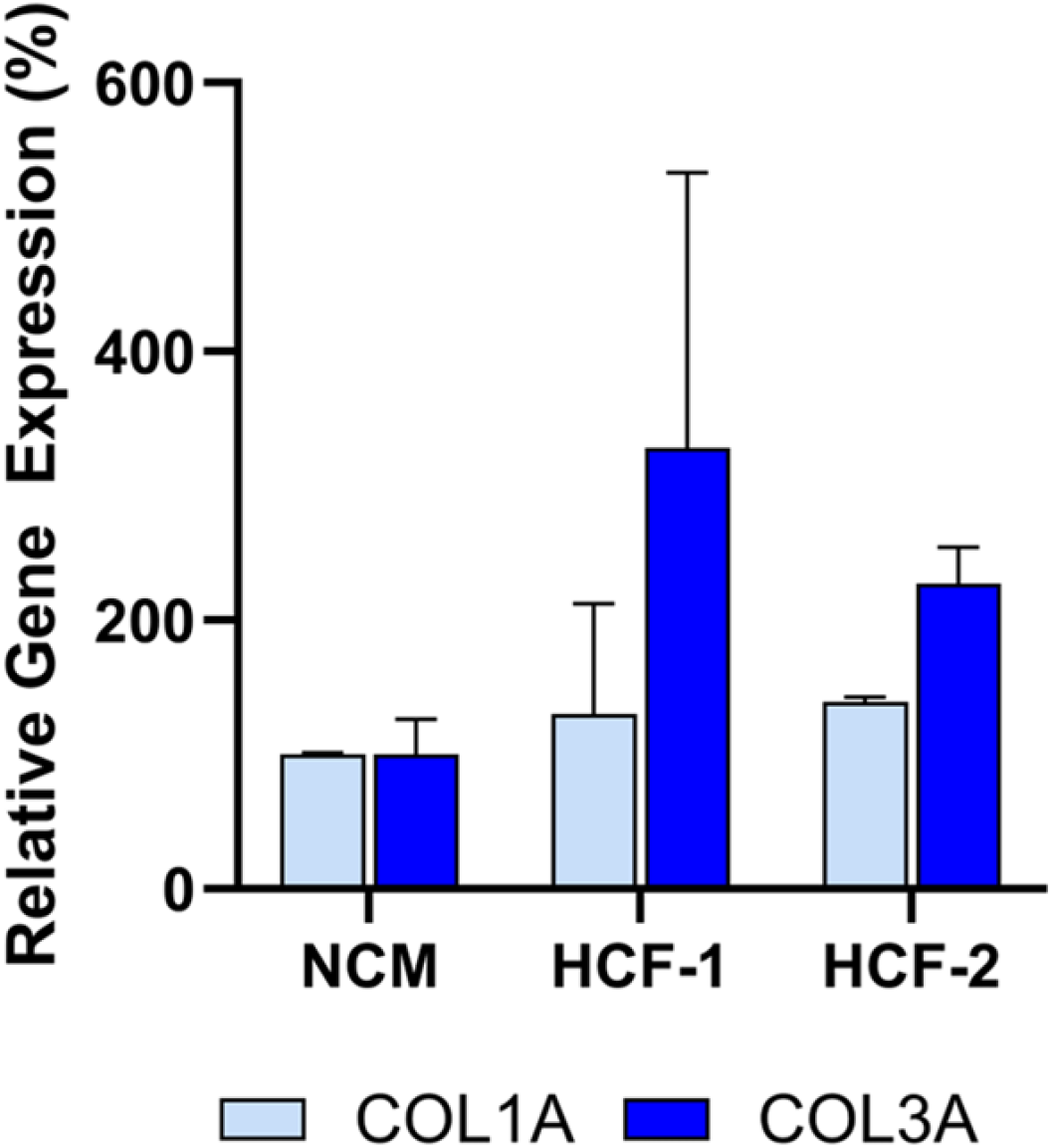
Human cardiac fibroblast (HCF) CM from two donors does not inhibit collagen gene expression (COL1A, COL3A) in fibroblasts after co-culture, demonstrating lack of anti-fibrotic activity. Assays were performed in triplicate, and data are presented as mean ± SD, normalized to NCM.

### 3.4. Dose-dependent Activity of Deramiocel CM in the Anti-fibrosis Assay

To demonstrate that the anti-fibrosis assay can detect changes in Deramiocel conditioned media (CM), CM from CDCs in two Deramiocel lots— LOT-0193 and LOT-0196—was serially diluted and evaluated in the assay. As shown in Figure 7, both lots exhibited dose-dependent inhibition of COL1A and COL3A expression. Non-diluted CM from both lots significantly suppressed collagen expression, confirming expected potency. Notably, increased dilution of both lots of CM resulted in less inhibition of COL1A and COL3A. At a CM dilution of 1:5 or higher, little inhibition of collagen was observed. These CMs would therefore be classified as non-potent. These results indicate that the assay is sensitive to changes in Deramiocel CM, such as reduced secretion of exosomes or growth factors, which are characteristic of less potent cells. Collectively, these findings support that the anti-fibrotic effect is mediated by secreted factors from CDCs in Deramiocel in a dose-dependent manner. For this study, recombinant human EGF (rh EGF) was used as a positive control for anti-fibrotic activity. The selection of rh EGF was guided by our preliminary studies using conditioned media (CM) from other cell types, such as human primary aortic endothelial cells (HAECs). When HDFs were treated with HAEC CM, we noted inhibition of COL1A and COL3A expression. Systematic component dilution experiments identified rh EGF as the mediator of collagen inhibition, leading to its adoption as a positive control to monitor if the assay is working properly (data not shown). Mechanistically, EGF engagement of the EGFR signaling pathway is known to modulate fibroblast activity and suppress excessive collagen production. Previous studies have demonstrated that EGF downregulates type I procollagen mRNA and protein levels in cultured human fibroblasts [22]. Other cardiac cell types were excluded due to the presence of EGF in their media formulations.

**Figure 7:**
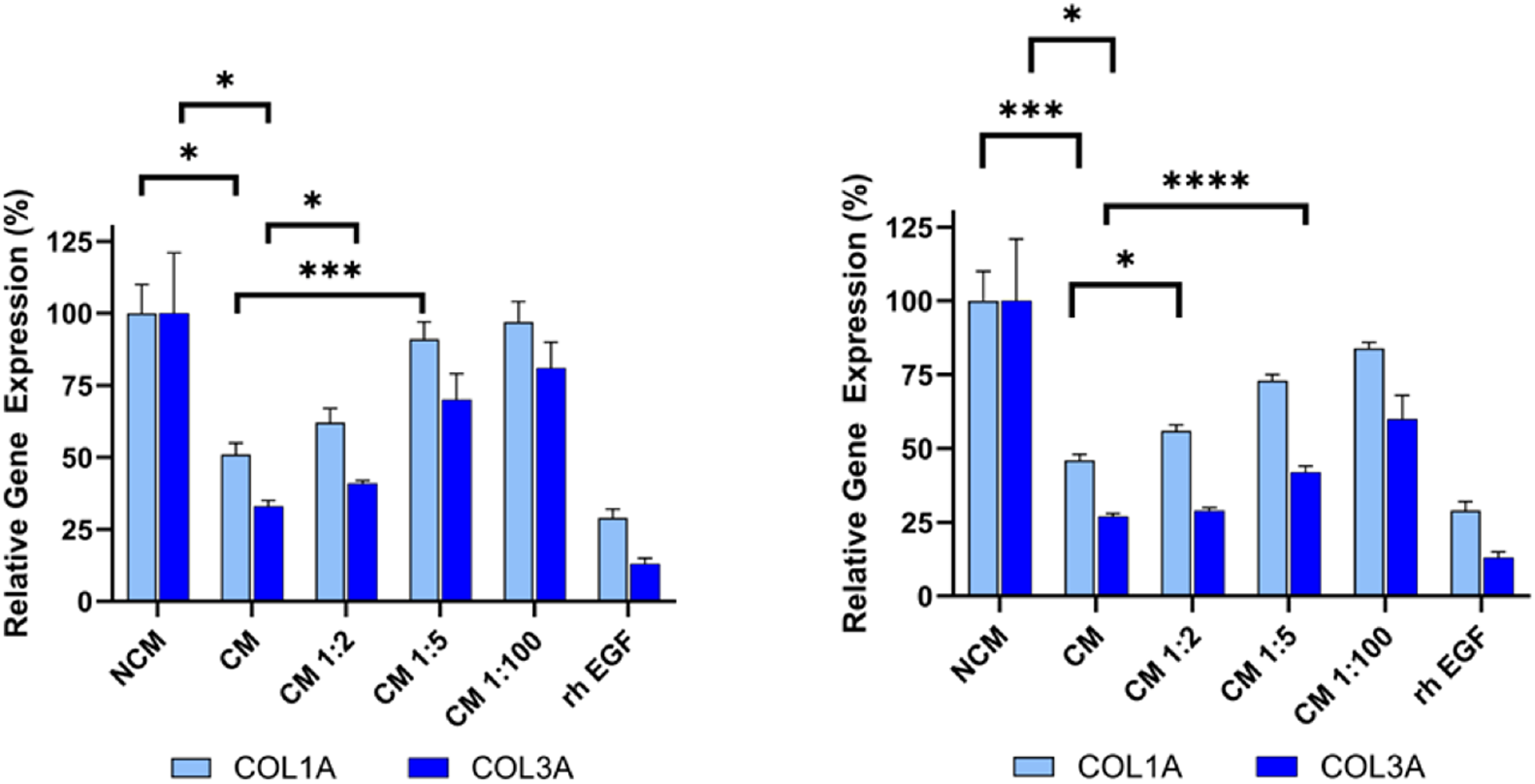
Dose-dependent inhibition of collagen gene expression (COL1A, COL3A) by serial dilutions of Deramiocel CM. Fibroblasts were treated with CM from LOT-0193 (left panel) and LOT-0196 (right panel). NCM and rh EGF (Positive) served as controls. Assays were performed in triplicate, and data are presented as mean ± SD, normalized to NCM. CM significance is marked against NCM and the first dilution with a significant p-value against 100% CM is marked **** p<0.0001, *** p<0.0005, * p<0.05.

### 3.5. Exosomes and Soluble Proteins are Responsible for Anti-fibrotic Activity

Deramiocel conditioned media (CM) contains two major components: 1) exosomes (and potentially other extracellular vesicles) released by CDCs in Deramiocel, and 2) soluble proteins, including growth factors secreted by CDCs in Deramiocel. Both components have been shown to contribute to Deramiocel’s mechanism of action [14, 21, 23]. Data presented above clearly demonstrate that Deramiocel CM, containing exosomes and growth factors, has anti-fibrotic activity.

To determine which components in CM mediate this effect, the role of exosomes in collagen inhibition was investigated first. CM from Deramiocel LOT-0193 and LOT-0196 was filtered using an Amicon 100 kDa column, which retains extracellular vesicles, including exosomes, ensuring the flow-through is exosome-free. The bioactivity of the exosome-free flow-through was then assessed in the anti-fibrosis assay. As shown in Figure 8, the rh EGF positive control significantly inhibited COL1A and COL3A expression, confirming assay functionality. Importantly, exosome-free CM exhibited increased COL3A expression compared to CM containing exosomes (i.e., CM, without Amicon treatment), suggesting that exosomes contribute to collagen inhibition (Deramiocel LOT-0196, initial data not shown, subsequent studies shown in Figure 8). These experiments were repeated three times across two lots, consistently showing increased COL3A expression following exosome depletion. Notably, COL1A expression was reduced to similar levels with both CM and exosome-free CM, indicating a differential effect of exosomes on collagen isoforms. These data strongly suggest that exosomes present in CM play a role in COL3A inhibition.

**Figure 8:**
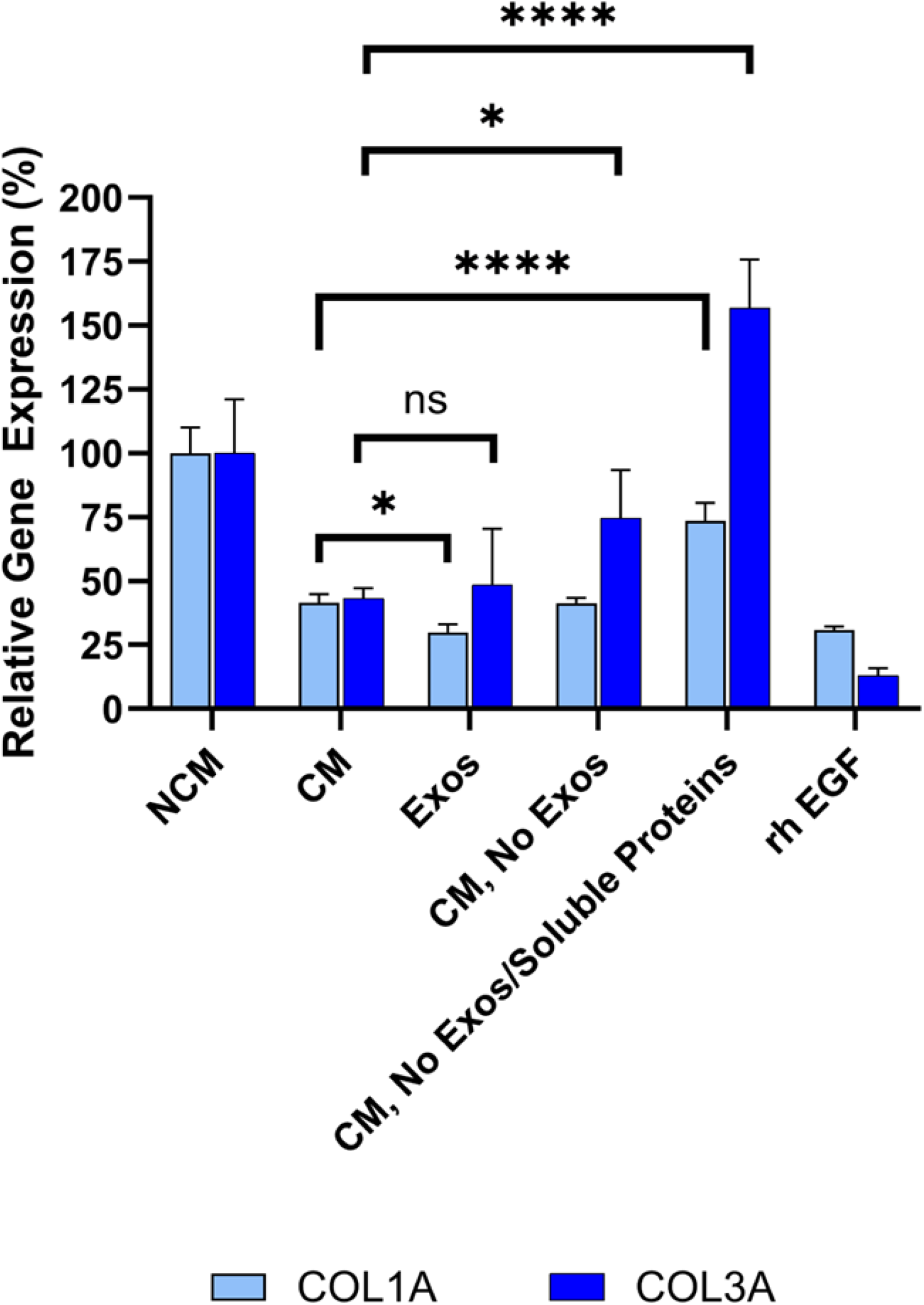
Sequential filtration demonstrates contributions of exosomes and soluble proteins to anti-fibrotic activity. Fibroblasts were treated with CM containing exosomes, exosome-free CM, or protein- and exosome-free CM, compared to rh EGF and NCM controls. COL1A and COL3A expressions were measured by qRT-PCR. Assays were performed in triplicate, and data are presented as mean *±* SD, normalized to NCM. **** *p <* 0.0001, * *p <* 0.05, ns = not significant.

To further dissect the contribution of soluble proteins in CDC CM in anti-fibrotic activity, CM from the same lots was sequentially filtered through an Amicon 100kDa column removing extracellular vesicles (e.g., exosomes), followed by a Amicon 10kDa column, which removes most soluble proteins (e.g., growth factors) from CM. Three fractions were collected from the sequential filtration: 1) exosomes retained in the 100 kDa filter, 2) exosome-free 100 kDa flow-through, and 3) exosome- and soluble protein-free 10 kDa flow-through. Bioactivity was assessed for each fraction alongside unfiltered CM. As expected, unfiltered CM significantly reduced COL1A (p<0.0001) and COL3A (p=0.0038) expression compared to non-conditioned media (Figure 8). Importantly, exosomes collected from the 100 kDa retentate induced a statistically significant reduction in both COL1A and COL3A expression compared to non-conditioned media. This reduction was not significantly different from CM, suggesting that CDC exosomes are necessary and sufficient for the bioactivity of CDCs, in alignment with in vivo studies.

Exosome-free CM (100 kDa flow-through) exhibited increased COL3A expression relative to CM containing exosomes, consistent with the prior depletion study (described above). Finally, exosome- and protein-free CM (10 kDa flow-through) demonstrated increased COL1A and COL3A expression to levels approaching or exceeding non-conditioned media, indicating loss of anti-fibrotic activity and classifying these samples as non-potent.

Collectively, these sequential filtration studies clearly demonstrate that both exosomes and soluble proteins, including growth factors, present in Deramiocel CM inhibit collagen expression and are responsible for the anti-fibrosis mechanism of action.

### 3.6. Secreted Proteins are Involved in Anti-fibrosis

Additionally, a multi-panel flow assay on deramiocel CM used in the anti-fibrosis assay revealed specific factors associated with anti-fibrosis, immunomodulation, and muscle survival. Similar to the anti-fibrosis assay, CM was compared with non-conditioned media (NCM) in the flow panel. Six factors with known direct or indirect roles in anti-fibrosis were observed to have an increased signal compared with NCM and are summarized in Table 1. Growth factors (FGF, EGF, and VEGF) are involved in anti-fibrosis via inhibition of myofibroblast activation, attenuation of the inflammatory fibrotic pathway and reduction of hypoxia-driven fibrosis, respectively and have been shown to be anti-fibrotic following injury [24, 25, 26, 27]. The cytokine IL-6 has been shown to be directly involved in the mechanism of action of CDCs by reducing myocardial scarring [28, 29]. The chemokine CXCL8 (IL-8) is known to be immunomodulatory and has been shown to be associated with improved left-ventricle ejection fraction (LVEF) following injury [30]. PECAM-1 inhibits endothelial to mesenchymal transition thus maintaining vasculature and reduces chronic inflammatory cell infiltration which both indirectly inhibit fibrosis [31]. These results, in conjunction with the filtration studies above, provide further evidence that the anti-fibrotic bioactivity of CDCs in deramiocel is mediated by exosomes and secreted soluble factors.

**Table 1:** Secreted factors known to play a role in the anti-fibrotic activity of Deramiocel Deramiocel CM.

| **Factor** | **Category** | **Role in Anti-fibrosis, Immunomodulation, or Muscle Survival** | **Supporting Reference(s)** |
| --- | --- | --- | --- |
| bFGF (FGF-2) | Growth factor | Potent anti-fibrotic; inhibits TGF-β-induced myofibroblast activation, preventing ECM production; promotes regeneration and tissue repair | [24-25] |
| EGF | Growth factor | Enhances tissue survival and repair, limiting scar formation; modulates immune cell recruitment, attenuating inflammatory fibrotic cascade | [26] |
| VEGF | Growth factor | Maintains vasculature and reduces hypoxia-driven fibrosis | [27] |
| IL-6 | Cytokine | Limits cardiac fibrosis; reduces myocardial scarring | [28-29] |
| CXCL8 (IL-8) | Chemokine | Modulates immune response in heart; prevents chronic inflammation; promotes cytokine release key to fibrotic regulation | [30] |
| PECAM-1 | Adhesion molecule | Reduces chronic inflammatory cell infiltration resulting in reduced fibrosis; role in anti-inflammatory signaling; supports vascular repair | [31] |

### 3.7. Potency and Efficacy of CDCs in Deramiocel Lots Utilized in Clinical Trials

Together, these findings demonstrate the anti-fibrotic activity of exosomes and soluble factors secreted by CDCs in deramiocel. The anti-fibrosis assay was subsequently qualified and validated as a potency assay to assess the bioactivity of CDCs in every lot Deramiocel for product release. Analysis of the anti-fibrotic activity of multiple clinical lots from the HOPE-2 trial and HOPE-2 open label extension study that have been shown to be clinical beneficial in DMD patients were used to set the acceptance criteria. Acceptance criteria for the anti-fibrosis potency assay correspond to <75% expression for COL1A and <72% expression for COL3A (equivalent to ≥ 25% and ≥ 28% inhibition, respectively), and were established based on historical lot performance across five working cell banks of reporter fibroblasts (data not shown).

The potency of Deramiocel lots administered in the HOPE-2 and HOPE-2 OLE studies was consistently observed in the anti-fibrosis assay with substantial reduction in expression of COL1A and COL3A observed in reporter fibroblasts after treatment with conditioned medium. Figure 9 presents expression values for the lots utilized in the HOPE-2 trial and HOPE-2 open label extension study. These trials demonstrated that DMD patients treated with Deramiocel experienced significant attenuation of disease progression, both in skeletal and muscle function. Importantly, long-term cardiac stability in DMD patients treated with Deramiocel was observed over 48 months in HOPE-2 OLE. This clinical effect, particularly in cardiac function, is attributable to the anti-fibrotic mechanism of action of CDCs as demonstrated in vitro by the anti-fibrosis potency assay. Thus the anti-fibrotic activity induced in vitro is consistent with clinical potency. The potency assay developed here was reviewed by FDA during regulatory interactions and was accepted as suitable for use as a potency assay for product release.

**Figure 9:**
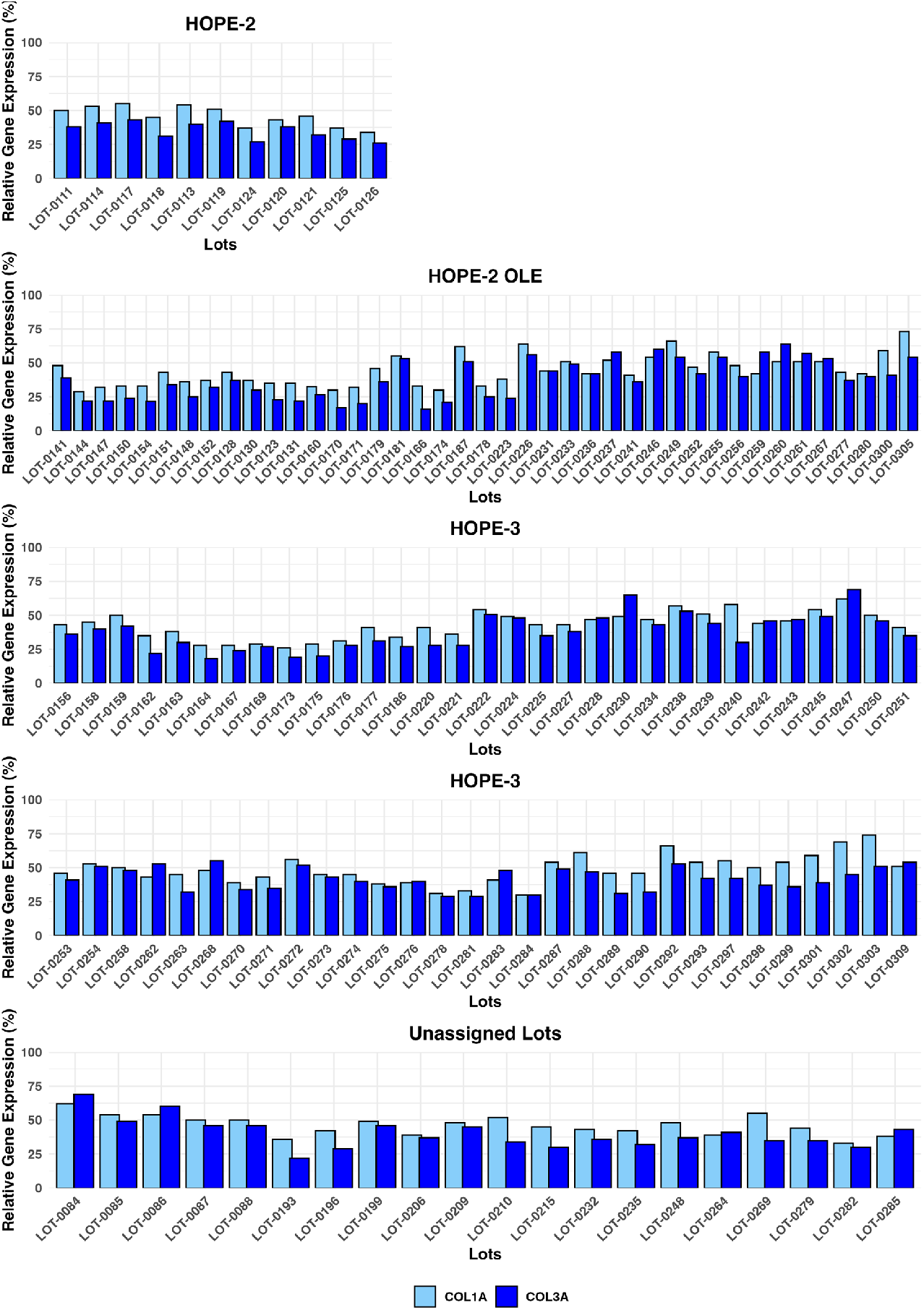
Deramiocel batches: COL1A and COL3A expression, measured in the anti-fibrotic potency assay. Data are shown for lots used in HOPE-2, HOPE-2 OLE, HOPE-3 (clinical data not yet unblinded) and lots unassigned to a trial, utilized for technology transfer or pending assignment to a clinical trial. Acceptance criteria are 75% for COL1A and 72% for COL3A (not shown).

In addition, data from the anti-fibrosis potency assay from all clinical deramiocel lots utilized in the ongoing HOPE-3 clinical trial, as well as lots that were not assigned to clinical trials, are shown in Figure 9. These data again demonstrate the consistent reduction in COL1 and COL3 by exosomes and soluble factors secreted by CDCs in deramiocel. Importantly, the inhibition of COL1 and COL3 met the acceptance criteria, deeming all lots as potent. All lots presented in Figure 9 were derived from master cell banks generated from >10 donor hearts, underscoring the reproducibility of the deramiocel manufacturing process. In total, conditioned media collected from CDCs in over 130 lots of deramocel were shown to induce a significant reduction in collagen when compared to non-conditioned media. The totality of this data demonstrate the potency of CDCs in deramiocel and the anti-fibrotic mechanism of action of CDCs observed in vitro, in vivo in pharmacodynamic studies and importantly in clinical trials.

## 4. Discussion

In this study, we demonstrate that conditioned media (CM) derived from CDCs in Deramiocel exerts robust anti-fibrotic activity in primary human fibroblasts. Across multiple independent lots, Deramiocel CM consistently suppressed expression of COL1A1 and COL3A1, two canonical markers of extracellular matrix deposition, while non-conditioned media or cardiac fibroblast-derived CM had no effect. These findings support the notion that the paracrine secretome of cardiosphere-derived cells (CDCs) harbors potent anti-fibrotic factors and provide mechanistic evidence for the therapeutic potential of Deramiocel.

Potency testing is defined as the ability of a product to achieve its intended therapeutic effect, demonstrated through appropriate laboratory assays (FDA guidelines and 21 CFR 600.3(s), 21 CFR 610.10). Regulations require that potency be assessed using in vitro or in vivo tests specifically designed for each product, ensuring a valid measure of therapeutic activity.

Initially, an in vivo myocardial infarction (MI) mouse model was utilized as a potency test for Deramiocel. MI is characterized by ischemic cardiomyocyte loss followed by adverse remodeling, including scar formation and ventricular stiffening. A major contributor to this remodeling is the excessive deposition of type I and type III collagens (COL1A and COL3A), which are key drivers of fibrosis and impaired cardiac function. Data from this model was used to assess and select deramiocel lots for use in early clinical studies and the completed HOPE-2 clinical trial. However, this model presents multiple limitations: it is labor-intensive, time-consuming (approximately several months per experiment), requires specialized surgical skills, relies on subjective echocardiographic measurements in small animals, and can yield variable, hard-to-validate results [32]. Moreover, the MI model does not replicate the progressive fibrosis characteristic of DMD-associated pathology [33, 34]. Indeed, mdx mouse models, the most common DMD model, demonstrate only mild, non-progressive skeletal muscle degeneration and do not faithfully recapitulate human disease, further reducing relevance for predicting clinical potency [35]. Therefore, development of a reliable and robust in vitro assay was required to assess the potency of each lot of deramiocel for product release.

Earlier work from the Marbán laboratory established that CDCs act largely through paracrine signaling rather than direct engraftment, with key mediators including epidermal growth factor (EGF), interleukin-6 (IL-6), and CXC chemokine ligand 8 (CXCL8/IL-8) [14, 36, 37]. In those studies, EGF secretion was linked to enhanced angiogenesis and cardioprotection, while IL-6 and CXCL8 contributed to immunomodulation and remodeling of the myocardial niche. Our results extend these insights by demonstrating a direct anti-fibrotic action of the Deramiocel secretome, as evidenced by downregulation of collagen gene expression in dermal fibroblasts. Importantly, we show that this activity is dose-dependent, exosome-associated, and robust across donor fibroblasts and Deramiocel manufacturing lots, thereby strengthening the translational relevance of these findings [14, 38]. Interestingly, we also detected high levels of IL6 and CXCL8 secretion in Lengendplex Flow Assay, which aligns well with the Deramiocel identity release assay using IL6 and CXCL8 as readouts. The assay measures these factors in RNA isolated from Deramiocel cell lysate by qPCR and demonstrates a strong fold-increase versus the negative control fibroblast RNA. EGF, measured by the Legendplex assay is also used as a positive control in the anti-fibrosis assay, further indicating its importance in the Deramiocel anti-fibrosis pathway.

The observation that CDC exosomes alone can recapitulate the anti-fibrotic activity of CDC CM and that exosome depletion abrogated the anti-fibrotic effects of Deramiocel CM highlights the central role of exosomes as vehicles for paracrine signaling [14, 36, 38, 39]. While prior reports emphasized soluble cytokines such as EGF [22], our data indicate that exosome-enclosed mediators may be the dominant contributors to the anti-fibrotic phenotype. Exosomes can encapsulate not only proteins such as IL-6 and CXCL8, but also microRNAs such as mi146a and mi148a [40, 41, 42, 43] and other regulatory cargos that suppress fibroblast activation and reduce collagen expression [14, 37, 44]. Thus, Deramiocel exosomes may represent a convergent delivery system integrating both cytokine signaling and gene regulatory programs.

The inclusion of fibroblasts from multiple donors in our study also provides a novel perspective, addressing biological variability often overlooked in earlier single-donor experiments. The reproducibility of Deramiocel’s anti-fibrotic effect across diverse fibroblast populations suggests that its paracrine activity is robust and donor-independent, an important consideration for clinical translation [36, 38]. Furthermore, the observation that cardiac fibroblast-derived CM failed to attenuate collagen expression underscores the uniqueness of the Deramiocel secretome relative to fibroblast paracrine signaling [38, 39].

Taken together, our work builds upon the foundation laid by the Marbán group by moving beyond descriptive profiling of CDC secreted factors to a functional demonstration of anti-fibrotic activity in primary fibroblasts. The innovation of our study lies in: 1) Establishing lot-to-lot consistency of Deramiocel CM in suppressing fibrosis-related gene expression. 2) Demonstrating that the effect is dose-dependent and exosome-associated, distinguishing Deramiocel from conventional soluble-factor therapy. 3) Providing evidence that cytokines previously linked to CDC function (EGF, IL-6, CXCL8) may participate in a broader anti-fibrotic signaling network when delivered via exosomes [14, 37, 38, 39, 44].

Importantly, the clinically meaningful improvements in cardiac and skeletal function observed in the HOPE-2 and HOPE-2-OLE clinical studies are directly attributable to the anti-fibrotic mechanism of action of CDCs in deramiocel. The anti-fibrotic activity of CDCs observed in the in vitro potency assay aligns with multiple in vivo pharmacodynamic studies in the mdx mouse model and large animal models, which demonstrate a decrease in fibrosis leading to a reduction in muscle scarring and consequently improved skeletal and cardiac muscle function. Importantly, the in vitro and in vivo anti-fibrotic mechanism of action of CDCs aligns with the therapeutic benefit observed in DMD patients after treatment with deramiocel. Thus, the validated cell-based anti-fibrosis assay employed herein offers a controlled and reproducible assay enabling a mechanistically informative assessment of Deramiocel’s potency and a demonstration of the deramiocel drug product to achieve its intended therapeutic effect.

## Author Contributions

Conceptualization, Yujia Li, Kristi Elliott and Minghao Sun; Data curation, Yujia Li and Marya Kozinova; Formal analysis, Yujia Li, Justin Nice, Marya Kozinova and Stephanie Adachi; Investigation, Yujia Li, Justin Nice and Stephanie Adachi; Methodology, Yujia Li, Justin Nice, Marya Kozinova, Kristi Elliott and Minghao Sun; Software, Marya Kozinova; Supervision, Yujia Li, Kristi Elliott and Minghao Sun; Writing – original draft, Yujia Li; Writing – review & editing, Yujia Li, Justin Nice, Marya Kozinova, Linda Marbán, Kristi Elliott and Minghao Sun.

## Declaration of generative AI and AI-assisted technologies in the writing process

During the preparation of this work the author(s) used ChatGPT in order to improve language and readability, with caution. After using this tool, the authors reviewed and edited the content as needed and take full responsibility for the content of the publication.

